# Mapping Mouse Brain Slice Sequence to a Reference Brain Without 3D Reconstruction

**DOI:** 10.1101/357475

**Authors:** Jing Xiong, Jing Ren, Liqun Luo, Mark Horowitz

## Abstract

Histological brain slices are widely used in neuroscience to study anatomical organization of neural circuits. Since data from many brains are collected, mapping the slices to a reference atlas is often the first step in interpreting results. Most existing methods rely on an initial reconstruction of the volume before registering it to a reference atlas. Because these slices are prone to distortion during sectioning process and often sectioned with nonstandard angles, reconstruction is challenging and often inaccurate. We propose a framework that maps each slice to its corresponding plane in the atlas to build a plane-wise mapping and then perform 2D nonrigid registration to build pixel-wise mapping. We use the L2 norm of the Histogram of Oriented Gradients (HOG) of two patches as the similarity metric for both steps, and a Markov Random Field formulation that incorporates tissue coherency to compute the nonrigid registration. To fix significantly distorted regions that are misshaped or much smaller than the control grids, we trained a context-aggregation network to segment and warp them to their corresponding regions with thin plate spline. We have shown that our method generates results comparable to an expert neuroscientist and is significantly better than reconstruction-first approaches.

## 1. Introduction

It is crucial to standardize and digitalize anatomical information to allow information from multiple brains to be used in a study and across different studies. To this end, detailed anatomical brain reference atlases have been established for both human and animal model studies (Hawrylycz et al., 2012; Lein et al., 2007). Ideally, all the experimental brain images would be automatically registered to an anatomical reference volume, creating a platform for the comparison and integration of different temporal and spatial experimental results. However, registering laboratory histological images to an atlas is still challenging in terms of accuracy, universality, and time-efficiency. One of the major issues is that brain histological data-sets often suffer from artifacts, such as enlarged ventricles (holes), missing tissue, folding, air bubbles, uneven staining, tears, and slice-independent distortions, shown in Figure 1. Existing programs mapping 2D histological sequence to a reference volume often require an initial reconstruction from these partially corrupted slices and therefore only work well with datasets of very good quality. However, histological datasets often require months of experiments to generate a result. Most labs still rely on manual brain region identification to fully utilize all the experimental dataset even if they are partially corrupted. This labor-intensive and time-consuming approach is highly variable and subjective among researchers.

**Figure 1:**
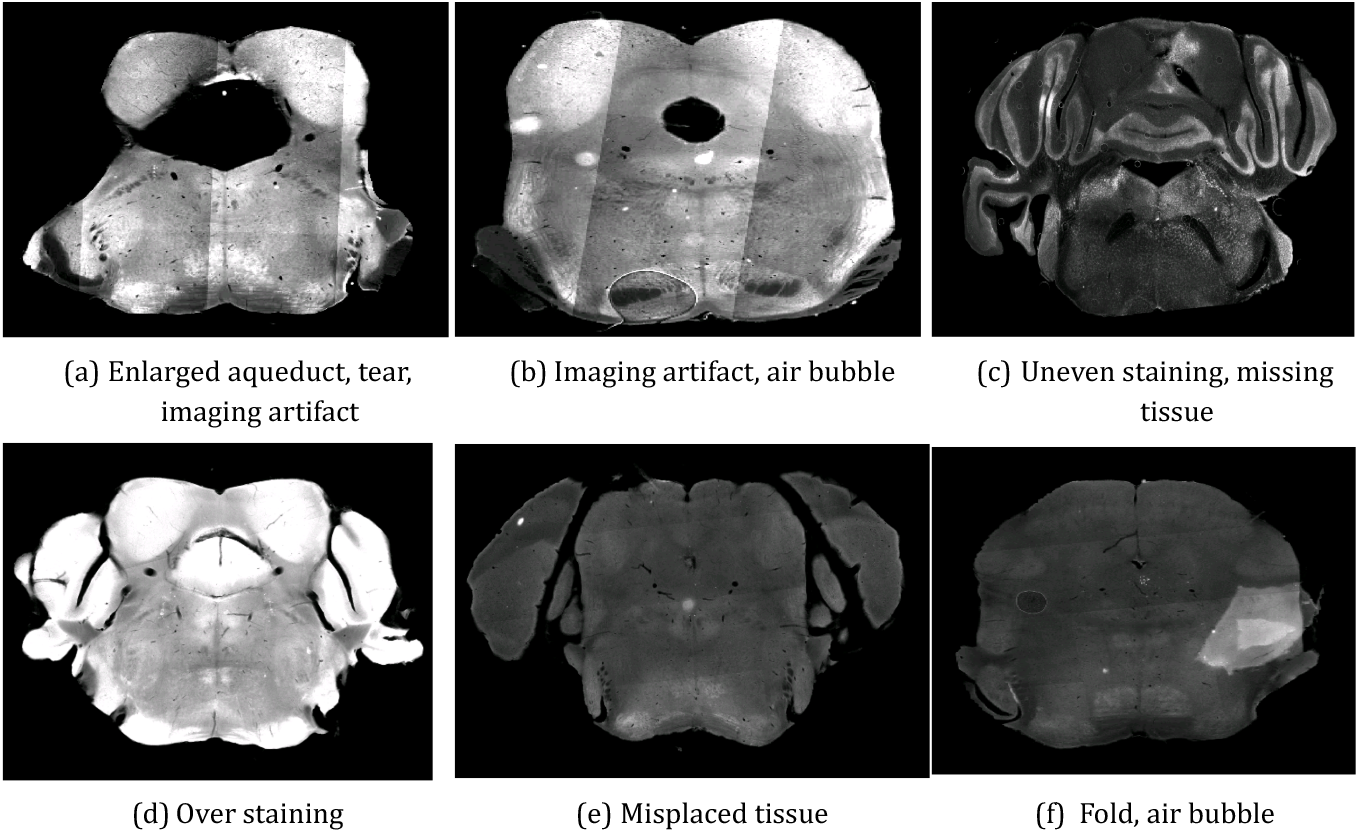
(a)-(f) Artifacts in histological brain slices. Images often suffer from multiple artifacts. (a) and (b) are contrast-enhanced for better visualization.

In this work, we introduce a method to register a sequence of coronal histological sections of mouse brain to the reference Allen Mouse Brain Atlas (Lein et al., 2007; all, 2015) by first identifying the matching sectioning plane in the atlas for each slice and then performing 2D nonrigid registration. The general idea and an example dataset is shown in Figure 2.

**Figure 2:**
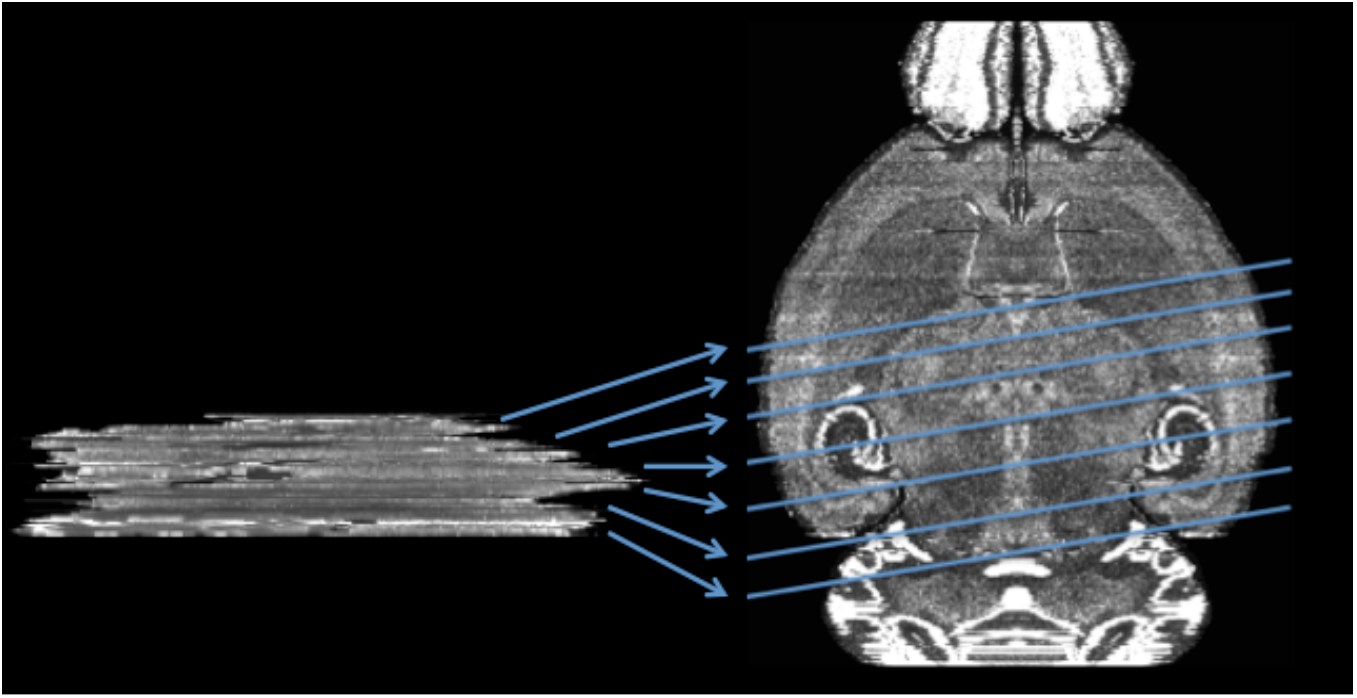
Mapping a sequence of histological mouse brain slices to the Allen Atlas (sagittal view). Left side shows a real histological stack. Right hand side is the Allen Mouse Brain Atlas. Atlas image credit: Allen Institute.

Our strategy makes the maximum use of the reference volume, successfully deals with the non-standard sectioning angle problem, preserves the curvature of the object - eliminating the z-shift problem (Adler et al., 2014), and is more tolerant to data corruption. This method takes into account some of the brain’s structural properties to minimize error, including the compressibility of different brain regions. The algorithm is tested both on full brain and sectional brain data, yielding faster and better correspondence than possible before.

## 2. Background

The problem of mapping a sequence of histological slices to a reference has been well studied. This prior work first reconstructs an initial volume estimate from the slices and then registers this reconstructed volume to the reference. Some works focus on the reconstruction problem because registration between a reconstructed volume and a reference is relatively standard (Stille et al., 2013; Dauguet et al., 2007; Mertzanidou et al., 2017); other works discuss on the reconstruction problem in an absence of reference volumes (Ourselin et al., 2001; Cifor et al., 2011; Ju et al., 2006; Bagci and Bai, 2010). Initial work reconstructed the experimental volume by pairwise registration of adjacent slices (Stille et al., 2013; Ourselin et al., 2001; Cifor et al., 2011). Due to tissue distortion, rigid registration is not sufficient. But pairwise nonrigid registration propagates any registration errors through the whole brain. This is especially problematic if any slice has a large deformation like missing tissue.

To improve overall reconstruction results and reduce error propagation, some methods align each slice with multiple neighboring images. For example, Ju *et al*. (Ju et al., 2006) reduced error propagation by warping each slice with a weighted linear and non-linear combination of warp fields to multiple adjacent slices. Others use blockface images (Dauguet et al., 2007) or select internal reference slices to reconstruct small chunks and then put together the entire volume (Bagci and Bai, 2010; Mertzanidou et al., 2017). However, with almost every slice at least slightly distorted, internal non-rigid registration will likely change the original shape of biological structures. Because this process tries to maximize the similarity between adjacent thin, *e.g*. 40 µm to 60 µm, histological slices, curved 3D structures along the sectioning direction may end up straightened. This 3D structure-straightening problem is known as the banana problem or z-shift (Adler et al., 2014). Once this error is introduced, it is hard to reverse completely even when this volume is registered to the reference. To avoid these volume distortion errors, one needs to use the reference volume earlier in the process by registering each experimental slice to its corresponding sectioning plane in the reference volume.

Recent work uses an iterative approach by first reconstructing a small volume and registering these slices to their corresponding planes in the reference (Yang et al., 2012; Goubran et al., 2013). Now the main challenge is finding the corresponding plane for each slice (Yang et al., 2012). This task is made more difficult because the experimental volumes have a non-standard sectioning angle, the brains are tilted in the sectioning machine, and have anisotropic resolution. The reconstructed volume has a very high resolution in the sectioning plane (determined by the resolution of the imaging system), and comparatively low resolution along the sectioning axis (limited by the minimum slice thickness). Yang *et al*.(Yang et al., 2012) selects a reference slice that maximizes the normalized mutual information after a 2D rigid registration is performed between a histological slice and each MRI slice. Goubran *et al*.(Goubran et al., 2013) registered each histological slice of a human brain to its corresponding MRI slice after the blocks are registered. While these methods work well when the sectioning angle difference is small, they introduce errors at larger sectioning angles.

To avoid these issues, we concurrently estimate the sectioning angle difference and the best matching planes in the reference atlas for each slice. This approach requires us to find the best matching slice in the reference before applying nonrigid deformations. Since the resulting slice comparisons are noisy, we aggregate information from all slices and use information about the brain’s structure to find the best match. Our method does not have a reconstruction step, therefore completely eliminating the z-shift problem. The details of our method are given in the next section.

After each matching reference slice has been determined, we need to perform a 2D registration between it and its matching histological slice. For 2D registration, Free Form Deformation (FFD) (Rueckert et al., 1999; Rohlfing and Maurer, 2003) has been the most common approach in neuroscience studies to map histological brain images(Jefferis et al., 2007; Geha et al., 2008; Dorocic et al., 2014; Costa et al., 2016). Mutual information is often used as the similarity metric because of image appearance difference caused by acquisition procedure variability. This method is highly dependent on the initial condition, because mutual information is known to be highly non-convex and has typically many local minima (Haber and Modersitzki, 2006). Because of staining variability within a slice, using mutual information does not always work. Instead, we found that the L2 norm of Histogram of Oriented Gradients (HOG) (Dalal and Triggs, 2005) difference suits histological slice properties better. Because HOG is non-differentiable, we base our work on the elegant discrete Markov Random Field (MRF) approach in (Glocker et al., 2008). Building on the annotation information of the reference, we build a MRF model based on tissue coherency. We further made improvement based on data-specific properties of our experimental dataset including segmenting a biological structure and warping them with thin plate spline (TPS) (Bookstein, 1989).

## 3. Method

This section describes in more detail how we find the sectioning angle difference and best matching plane in the reference volume for each histological slice (Section 3.1), and nonrigidly register each slice to the corresponding sectioning plane in the atlas slice (Section 3.2).

Both in the 2D to 3D localization and the 2D nonrigid registration steps, a relatively sensitive and quantitative similarity measure is needed. The state of art is to use normalized mutual information (Jefferis et al., 2007; Geha et al., 2008; Dorocic et al., 2014; Costa et al., 2016). Despite its wide use, it didn’t work well in our images since this metric fails when intra-slice uneven staining causes intensity variability within a structure which breaks the statistical correlation between a slice and its target image.

When searching for a better metric, we also wanted to find one that would work well for our images. Our image characteristics include:

1. Staining reagent and microscopic setting difference can cause direct comparison of intensities to be not useful. Even worse, due to non-uniformly applied staining reagent, some slices are unevenly illuminated.
2. NISSL-stained (Glaser and Van der Loos, 1981) images which only highlight the cell body of neurons. Two matching images will show corresponding anatomical structures but do not have pixel-wise cell body level correspondence.
3. Sparsely scattered or densely populated cell bodies which make images low-contrast and noisy. Many descriptors that work with man-made scenes do not perform well.
4. Distortions caused by brain’s elasticity requiring metrics that work even when the two images are slightly distorted from each other. This distortion tolerance also allows it to compare a distorted histological slice to a reference slice.

As shown by Dalal in (Dalal and Triggs, 2005), HOG has the capability to deal with pose, illumination, and background variations which mimic many of the issues in our images. We adopt HOG and use the L2 norm of HOG difference between two images as the similarity metric. To use this metric the two images first are brought to the same coordinates with a similarity transformation estimated with the Umeyama method (Umeyama, 1991) based on contour point correspondence generated by Shape Context (Belongie et al., 2000). Smooth tissue contours are extracted by applying the Fourier transform on the boundary curve and removing high frequency components. To accommodate the global deformation caused by the force in the direction of sectioning, slices are further rescaled in the horizontal and vertical direction. We then extract HOG features from both images and generate the scalar result by summing the squared difference between HOG feature vectors for each block at the same coordinate in the two images. For 2D to 3D localization, a large cell size is used to be less sensitive to local distortion. For 2D registration, we find nonrigid transformation to minimize HOG L2 norm with a small cell size.

### 3.1. 2D to 3D Localization

Since histological slices are often cut with near parallel angles with a microtome, it is fair to assume parallel cutting angle throughout the whole brain. Because atlas is uniform in each dimension, to find the cutting angle difference, we rotate the atlas with different angles, resection it into coronal slices, re-index the slices in order, and compare the new resectioned atlas slices to the histological sequence.

The following sections give our dynamic programming formulation to solve the alignment problem to determine the slicing angle, and a simple method to increase sensitivity to angular shifts.

#### 3.1.1. Slice Mapping with Dynamic Programming

The best cutting angle is the angle that maximizes similarity between all histological slices and their corresponding best matching slice in the atlas. Because in-plane rotation can be fixed, we only consider rotation angle *α* about the superior-interior (y) axis and *β* about the left-right (x) axis. To solve the problem, we first find the best matching slice for each experimental slice given a potential cutting angle. The problem can be represented as follows: Let I_1…N_ with spacing s_E_ be the experimental slice sequence, and *V_A_* be an isotropic reference atlas with voxel dimension s_A_, defined on the domain Ω. After rotating the atlas with potential best rotation *R_αβ_*, we reslice the rotated atlas into coronal slices and re-index them as atlas slice sequence J_1…M_. Using the L2 norm of HOG differences described in Section 3, we aim to find a mapping that matches each slice in I to a slice in J which minimizes the overall difference.

Taking into account potential compression along the longitudinal axis, slice quality variation, and inter subject variation, we formulate the problem with a single subset A of all slices I, where A is an ordered selection of 1… *N*, which may or may not be the whole sequence of experimental slices (depending on the image sequence quality and the value of s_A_ and s_E_). A is chosen to span the full sequence while avoiding damaged slices.

We formulate this slice mapping and difference minimization problem as a dynamic program. Let I_A_ be the ordered selection of experimental slices, and let J be the resliced atlas sequence ordered from the same direction along the longitudinal axis and spacing sA. The cost, *C*(*i, j*), is defined as the minimum cost of mapping the first i slices in A to a sequence of j slices, where the *i^th^* slice has to be mapped to the *j^th^* slice:

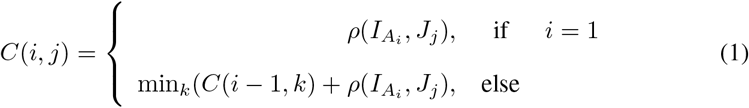

where *i ∈ A*, 0 *≤ j ≤* **card**( *J*), *ρ*(*a, b*) denotes the difference score between Slice a and Slice b measured with the HOG similarity metric. To reduce the required computation, we only look at potential matching slices, k, which have plausible spacing:

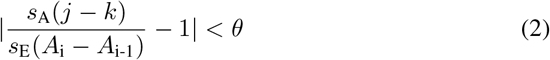

where *A_i_* is the original index of the *i^th^* slice in the selected sequence and *θ* is a userdefined threshold value. This spatial constraint constrains the ratio of the distance between slices in the atlas and the experimental slices match to *θ*.

We denote the best *k* that satisfies Equation 2 and is used to fill in the cost matrix (Equation 1) as *k^∗^*. The best intermediate steps are saved by updating the three-dimensional array *M* for each *i, j*:

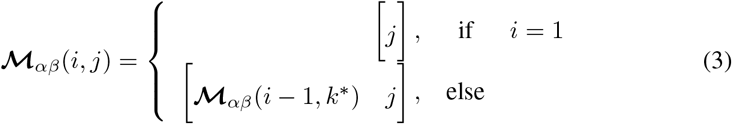

***M****_αβ_* (*i, j*) lists the the indices in J that best match each of the first i slices in A, where Slice i in A is mapped to Slice j in J and atlas is rotated with angle *α* about y axis and *β* about x axis. The optimal mapping is therefore given by:

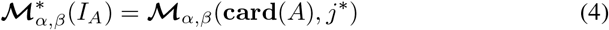

where

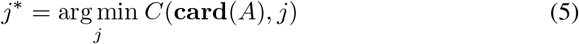

The cost of mapping all slices in A to resectioned slices in J with atlas rotated by *αβ* is given by *C* (**card**(*A*)*, j^∗^*).

#### 3.1.2. Cutting Angle Difference Determination

After running this dynamic program with different sectioning angles we could directly choose an angle that gives minimum cost score to be the best cutting angle. However, since HOG is relatively insensitive to local distortion and each slice is slightly distorted, when summing up all the costs we are also summing up a lot of noise. Therefore when the angle is very close to the true sectioning angle, the difference among neighboring angles is not substantial. To improve our robustness, we use a different approach for angle determination. This approach also predicts how we should adjust the rotation preventing the exhaustive search required in the previous approach.

Biological structures change quickly along the posterior-anterior direction. It is not hard to tell if an experimental brain is sectioned with a different angle than a reference atlas, even if the angle deviation is only several degrees, because structures that appear in the same slice in the atlas will be in different slices in the experimental slice sequence. For example, if the left side of a brain is tilted to be more anterior, on average the right hand half coronal brain slice will appear to be more posterior to the left half. Thus if we match the left and right half slices of an experimental brain separately to the atlas, we will see that the slice number of the matching slices of left half brain will be on average higher than that of the right half brain. Based on this idea, we use matching slice index difference of half brains to determine if a rotation angle best fixes the cutting angle difference between the experimental brain and the atlas.

Because mouse brains are generally symmetric along the superior-interior (y) axis, the rotation angle *α* about this axis tends to be around zero. The rotation angle *β* about the left-right (x) axis tends to be larger because mouse brain is not flat at the bottom and can easily be set tilted on the microtome plate. Here we use the determination of angle *β* as an example; the flowchart is shown in Figure 3.

**Figure 3:**
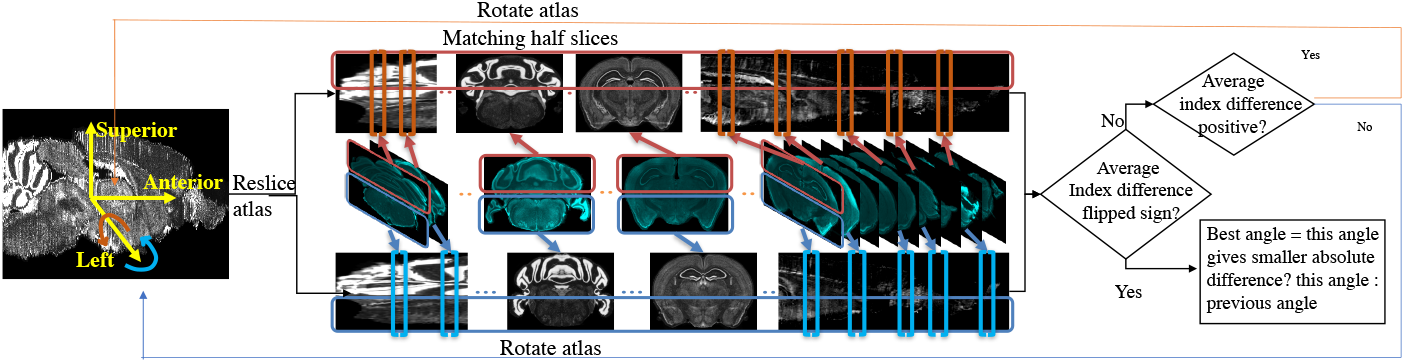
Flow chart for determining sectioning angle about the left-right (x) axis. In the matching half slices step, atlas is stretched for better illustration. Atlas image credit: Allen Institute.

To find the best rotation angle *β* about x axis, we solve the slice mapping problem with method described in Section 3.1.1 on the upper half slices and the lower half slices respectively. We take the index difference between the optimal mapping given by:

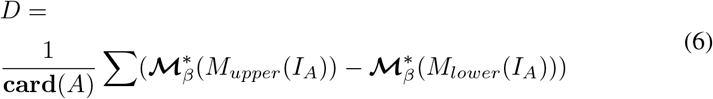

where M_upper_ and M_lower_ denotes binary masks to apply to both experimental image and resliced atlas image. Positive *D* means upper half experimental slices are matched to slices more anterior to those lower half slices are matched to. Therefore the atlas should be rotated more in postive direction about the left-right (x) axis, where the positive direction is defined by the right-hand rule around the positive direction of the x axis. If *D* is negative, then the atlas should be rotated in the negative direction. We reslice the atlas again after rotation, and repeat the above steps until the index difference flips sign meaning we need to rotate the atlas with another direction. The rotation angle change has step size of one degree. When the flipping of sign occurs, we choose the angle between the current angle or the previous angle whichever gives the smaller absolute index difference. The same steps are repeated to determine *α*.

After finding the optimal rotation 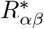, we apply the mapping method on full slices in A to find their corresponding full slices in the optimally rotated atlas. We then interpolate linearly on the matching slice indices to find the best matching slice for every other experimental slice in the experimental volume that is not selected in A.

### 3.2. Coherency-based 2D Deformable Registration

After the 2D to 3D registration, we register all the experimental slices nonrigidly to their computed corresponding slice in the optimally rotated atlas to build a pixel-wise mapping from the 2D slice sequence to the reference volume. Let an experimental image g that is globally transformed to the same coordinates of its corresponding slice be the target image, and its best matching slice f be the source image, where Ω ⊂ ℤ^2^ is the image domain. In the task of 2D registration, we aim at finding a transformation *T* such that

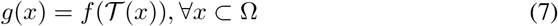

where g and f become equivalent in terms of anatomical structures.

For registering histological images, the most common approach has been mutual information based Free Form Deformation (FFD). Like in FFD(Rueckert et al., 1999), we superimpose a uniformly spaced sparse grid *𝒢 ⊂* Ω. Because of the properties of the experimental images described in the previous sections, we continue to use HOG difference as the similarity metric but with a smaller HOG cell size to fix local distortions. Because HOG is not differentiable, we build our work on a discrete Markov Random Field (MRF) approach (Glocker et al., 2008), where for each node *p* ∈ *𝒢* we seek to assign a label *l_p_* ∈ *L* that minimize an energy function *E* consisting of a unary term that ensures similarity - the HOG term in our case - and a pairwise term that regularizes motion between neighboring nodes. Each label *l^p^* corresponds to a displacement *d* in a predefined displacement set Θ by which we displace a node. We define the bijective function b between L and Θ as *b*: *d → l*.

#### 3.2.1. Model Elasticity with the Pairwise Term

The ventricular system spans throughout the brain, providing fluid pathways in the brain, and creating regions of “empty” space in almost every histological slice. Those cavities are easily deformed during sectioning procedures and have much inter-subject variation. Thus when computing this MRF warp field, one needs to take into account the elasticity of different regions in the brain. By warping images to match with each other, we are essentially warping tissues: the more two adjacent control points are displaced the more tension accumulates, if the two control points are connected through coherent tissue. In contrast, if they are separated by any empty space, no tension should be built in between.

The traditional and most common interpolation method for biomedical image analysis has been the B-spline model (Rueckert et al., 1999), where each pixel is affected by 4 × 4 neighboring control points. In the case where two control points are separated by an empty space, a B-spline interpolation no longer makes sense because of discontinued tissue coherency. Therefore to better model tissue deformation, we use the simple bilinear model where a point is only affected by its direct 2 × 2 neighboring control points. Of course now our warp generation needs to ensure some smoothness.

Our system does so with a very simplistic model. We divide each target slice into two regions: free (ventricular system and background), and coherent (other areas) based on atlas annotation. We then classify the nodes as coherent (red) or free (green in Figure 4) based on if they are inside a coherent or a free region. The idea is tension only accumulates when we compress or stretch two nodes that are connected solely with coherent region. If there is an empty space between two nodes, intuitively compressing them or stretching them should not build tension in between. Based on this property, we use the pairwise term - the traditional regularization term - to model tension accumulated between nodes with which we stretch or compress brain tissues.

**Figure 4:**
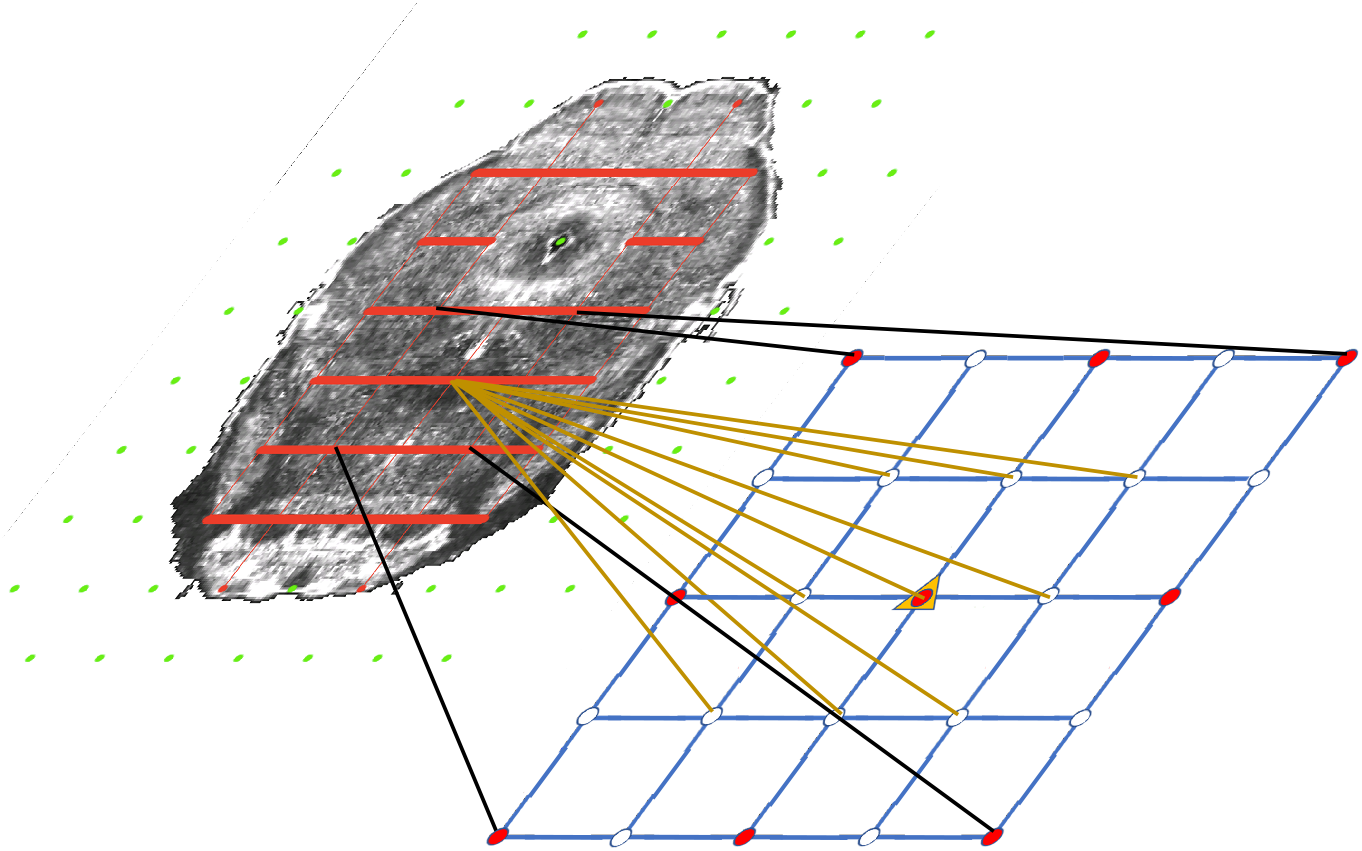
Illustration of the coherency model and grid refinement from level t to level t+1. Grids are overlaid on an atlas image (contrast adjusted for better illustration) to show the coherency model. Green nodes are free nodes which include nodes in ventricle systems and near the boundary of the tissue in the background. Red nodes are coherent nodes which can’t be seen since they become part of the tension edges, represented by red line segments between coherent nodes. During refinement, the grid size is decreased by two in each direction quadrupling the number of nodes, which is shown in the lower grid. The motion of existing nodes are carried onto the next level. The motion of non-existing nodes in the lower grids are interpolated from the motion of the existing nodes.

There is an edge (red line segment in Figure 4) only if the connecting line segment between the two nodes only crosses coherent regions which indicates both nodes must be coherent as well.

We first extract mask images *r_c_* and *r_e_* representing coherent region - tissue - and empty space region - ventricles and background - respectively from the reference annotation and project them to the source image. We further group control points as coherent or free in Eq. 8. Coherent control points are inside coherent regions. Free control points are the control points inside an empty space, and moving the control point will affect pixels inside coherent regions.

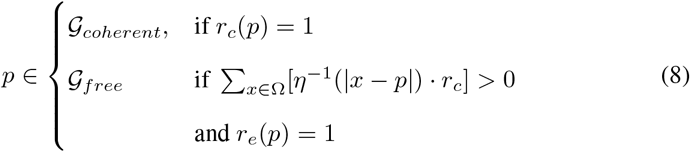

where the inverse influence function, *η*(.)^−1^, adopted in (Glocker et al., 2008), masks pixels influenced by a control point p. We include the influence function in control points classification, because we only care about control points that affect image appearance.

We further define a tension edge set, **E**, where tension accumulates when moving the two control points connected by an edge in this set. Basically an edge *e_pq_* is in **E** if the line segment connecting Node *p* and Node *q* only crosses coherent region *r_c_*. Because the spring potential energy is proportional to the square of displacement, we use squared difference as the pairwise term:

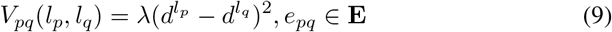

where *λ* is a regulation parameter.

#### 3.2.2. Multi-level Estimation

We need to be able to both correct large distortion, and make small changes to achieve good results. For both computation efficiency and quality of results we chose to use a multilevel approach. Since we are trying to model the tension that the deformations create, we need the pairwise energy terms to accumulate as we refine the grids. This requirement means we can’t use the approach used by Gloker *et al*. (Glocker et al., 2008), and instead solve the problem using a method where each refinement level maintains knowledge of the distortions created by previous levels.

The conventional multilevel approach (Glocker et al., 2008) repeats the same procedure with progressively finer grids: locations of the control grid points are computed that minimize the unary and pairwise term, and then the resulting image is warped to match these new grid locations. The next level grid is added to the warped image and the process is repeated. To maintain tension in a realistic way, we do not reset the grids and tension after each iteration, and use each iteration to simply update the allowable possible positions (labels) for the next iteration. More formally to carry the squared form tension correctly to the next level, we expand Θ to contain all possible discrete displacement of a node over all iterations and represent possible displacement at level t to be *D_t_*. At the same time, we expand 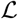 so that bijective relation *b*: *d → l* still exists.

While in this multilevel approach, the pairwise term is straightforward and without any change, estimating the unary term measuring similarity is more complicated. As mentioned above, we first need to compute the set of possible locations for each grid point, which depends on the results from the prior level. To do this, we denote the grid at level t as *𝒢_t_* and the influence function as *η_t_*. Let 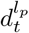 be displacement at node *p* with label *l* at level t. At each level t, we estimate best motion 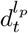 at each node *p* ∈ *𝒢_t_* and bilinearly interpolate them to get the initial displacement for each node in *𝒢_t+1_* at the next level. We denote this preset displacement at node *p* as 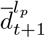, therefore the set of possible displacement at node p level t+1 is given by the sum of this preset displacement and possible displacement in level t+1:

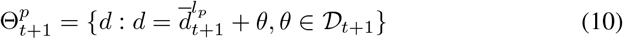

Having created the set of possible locations for each of the grid points, we next need to create the image that we will compare at this level for the similarity measure. In previous work, this warped image is input to this level, but we need to compute that image from the displacements of the previous level’s control points and the labels associated with the node we are evaluating. When estimating the local patch around node *p* at level *t* + 1, we use the positions of the grid points that are around p from the prior level and the position of *p* for the given label. This provides an estimate that incorporates the warp of the prior level and an estimate of the additional warp created by moving p to the position indicated by this label. For simplicity and computational efficiency, this estimate ignores the warp that will be caused when other control points at this level move.

We denote the patch that is affected by p in the first level function 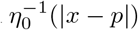 as 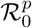. The control points in the patch at level t+1 is defined as:

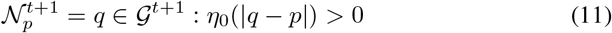

To create the image that we will compare, we set the nodes in *N_p_* at the values from level *t*, except for node *p* which we set with the displacement for the current label in the set 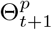. Therefore the transformation applied to the affected region 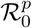 when we associate label *l_p_* with node *p* at level *t* + 1 is:

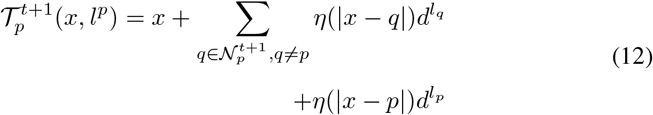

Thus the unary term is given by the similarity measure between the warped patch and target patch:

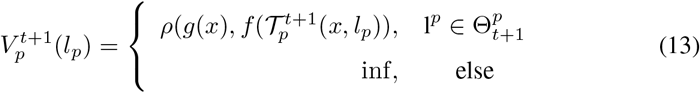

where 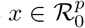 and *ρ* measures the difference score between two images.

Eventually we formulate the MRF energy function at level t as the summation of the normalized unary term and pairwise term:

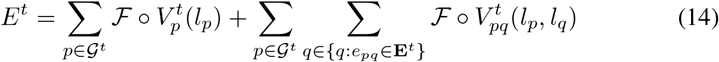

where *𝓕* denotes the normalization operation. Because each patch has various apear-ance, we feature scale them so that each node contributes equally and is within range [0, 1]. Because the free nodes are not constrained with any pairwise term, they are essentially assigned labels that minimize the unary term:

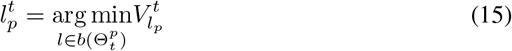

where 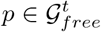. Coherent labels are solved by minimizing the energy function:

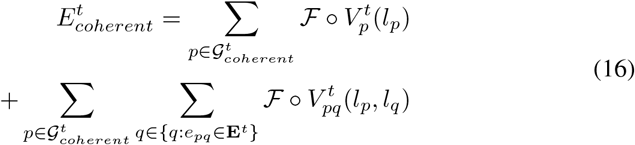

This tissue coherency based model enables us to generate close to human level results as described in Section 4.4.

#### 3.2.3. Dealing with Unremoved Background in the Atlas

The Allen Mouse Brain Atlas has very low intensity pixels around real brain tissues as shown in Figure 5. Minimizing HOG difference on the free nodes that influence tissue contours will work if both the target and source images are preprocessed to include only the real tissue. However because of the background noise in atlas images, and HOG’s relative insensitivity to intensity, this noise can cause significant errors in contour alignment. We added an intensity threshold in the process of computing HOG - if a pixel’s intensity is lower than the threshold, its gradient is not included in the histogram. However this does not solve the problem because a single threshold intensity cannot eliminate the background noise perfectly.

**Figure 5:**
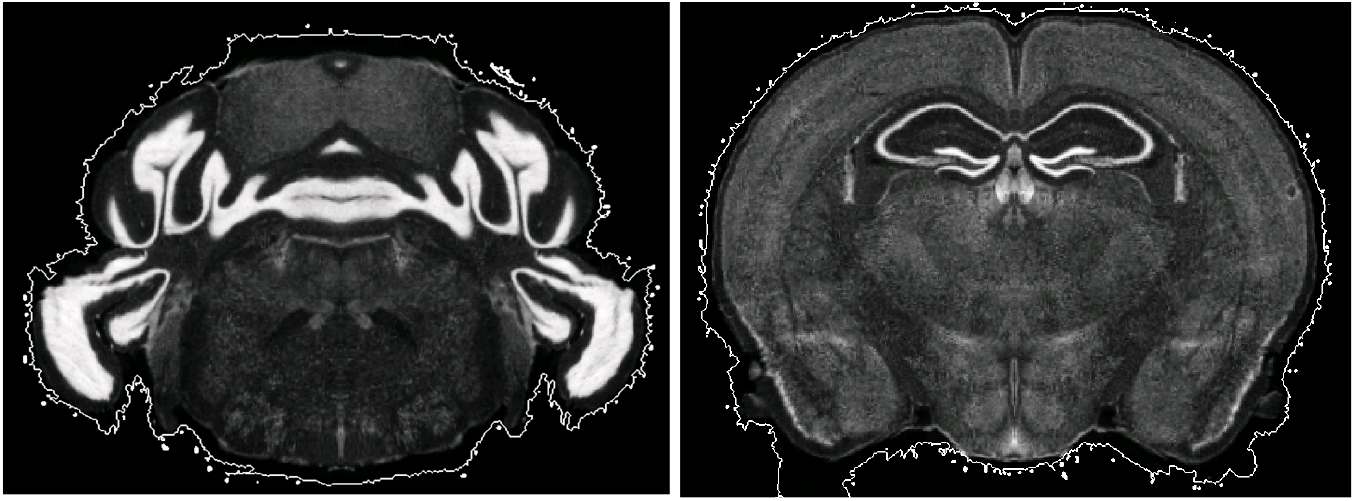
Example atlas slices with nonzero intensity regions circled by white contours.

Because we are trying to align the contours, we take an alternative approach which is more robust to this noise. Since the extracted contours won’t be perfect, we smooth them, and then align them to minimize the non-overlapping regions. The contours are generated by 1) thresholding the original image, 2) taking the largest binary region, 3) filling holes in the largest binary region, 4) morphological eroding and dilating so that the masks are smooth.

We mask the experiment image contour pixels by *c_e_*, and real tissue in image *f* and *g* to be *m_f_* and *m_g_*. If a node *p* influences any contour pixels, we modify its unary term to include an additional cost related to the number of unmatched pixels:

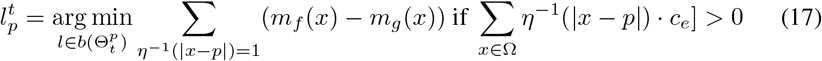

### 3.3. Improvement based on Data-specific Properties

Our framework was used in a systematic anatomical study in the hind brain to map the brain regions containing the dorsal raphe nuclei to the Allen Mouse Brain Atlas to study organization of the dorsal raphe serotonin system and its behavioral functions related to depression and anxiety (Ren et al., 2018) - attached as an appendix. The dorsal raphe nuclei is ventral to an empty space called aqueduct. Due to difference in brain preparation procedure, the aqueduct in some slices are highly misshaped as shown in Figure 6.

**Figure 6:**
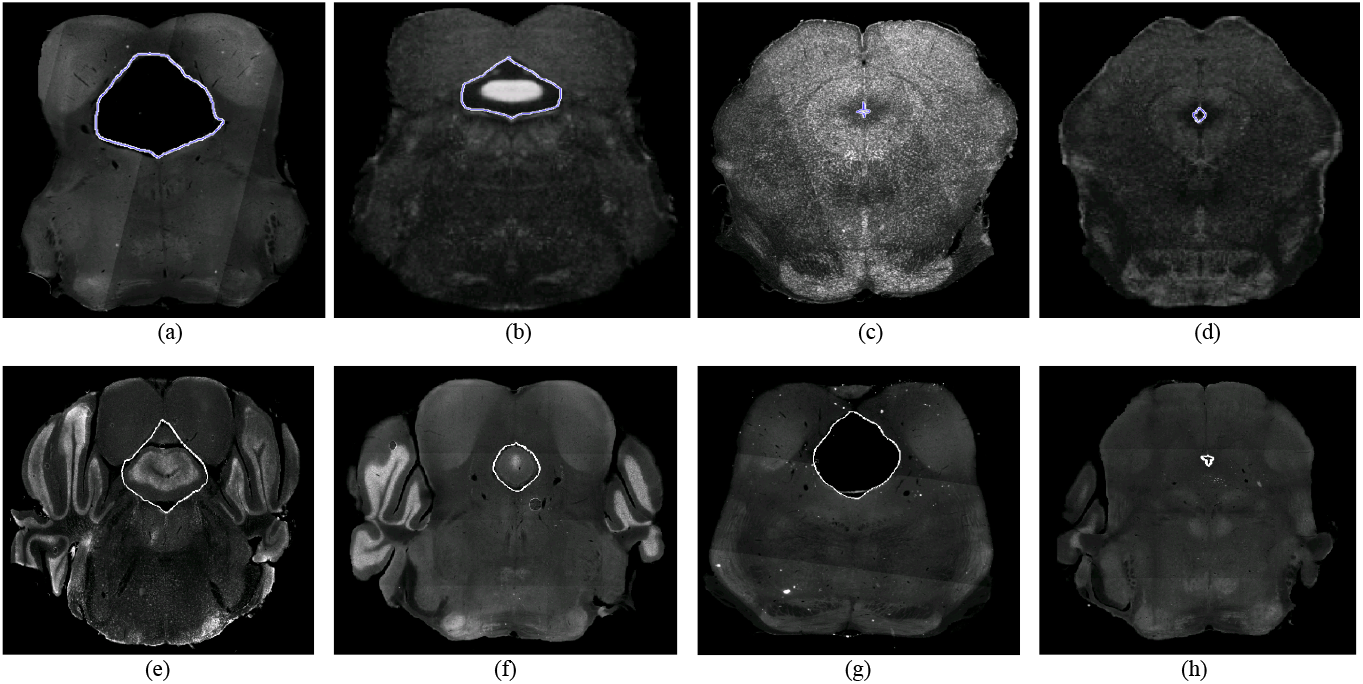
(a) Expanded aqueduct in an experimental image. (b) Normal aqueduct in the corresponding atlas image of (a). (c) Squeezed aqueduct in an experimental image. (d) Normal aqueduct in the corresponding atlas image of (c). Aqueduct contours are labeled in blue. (e)-(h) Various aqueduct appearance in different brains and different slices. Aqueduct contours are marked with white curve.

The significant difference in size, appearance, and edge orientation of the aqueduct makes aligning the regions around it difficult using the coarser grained HOG descriptor alone. This situation is made even more difficult because a squeezed aqueduct can be smaller than the grid size in the finest iteration.

We solve the squeezed aqueduct problem by warping the segmented aqueduct to the corresponding annotated atlas aqueduct with TPS. Because aqueduct appearance varies across subjects, sectioning angles, and longitudinal axis as shown in Figure 6, it is hard to segment them with a single traditional segmentation method. We trained manually labeled aqueduct with a network similar to Chen *et al*.’s (Chen et al., 2017) implementation of context aggregation (Yu and Koltun, 2015), a convolutional network designed for dense prediction. To build point correspondence on the aqueduct contours, we find the highest and lowest point on the aqueduct contours. If there are multiple highest or lowest points, we choose the point that is closer to the centroid of the aqueduct contour. We divide the contour in halves with the highest and lowest points and build point correspondence by uniformly sampling the same number of points along the curve. The points on the aqueduct contours can only ensure the alignment of inside the aqueducts. Since the images are mostly aligned with MRF, we include the control points outside of both aqueducts to add more control to the tps warp.

With point correspondence on the aqueducts’ contours, we add another term to the unary term so that the aqueduct are brought closer before reshaped with TPS. The term measures Euclidean distance between the warped experimental aqueduct contour points and their corresponding atlas aqueduct contour points:

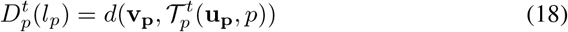

where an experimental aqueduct contour point *u* ∈ **u_p_**, if its influence to node p - η_t+1_ (*|u − p|*)>0. **v_p_** are the corresponding aqueduct contour points in the atlas image, and *d* measures the Euclidean distance between two sets of points. Therefore the energy function for coherent nodes in Eq. 3.2.2 becomes:

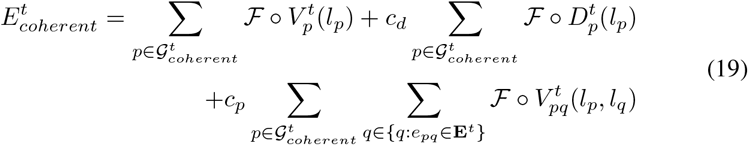

where *c_d_* and *c_p_* are the coefficients before the point distance unary term and the pairwise term respectively. We assign labels that minimizes the new combined unary term to the free nodes:

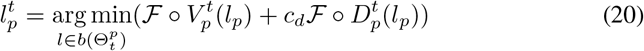

where 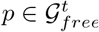.

## 4. Experiment

Our framework was developed to register a full mouse brain slice sequence consisting of 202 60 µm thick slices to the Allen Mouse Brain Atlas and was also used in a systematic anatomical study in the hind brain to study the organization of the dorsal raphe serotonin system. We successfully mapped all 36 sectional mouse brains in this study to the Allen Mouse Brain Atlas. Each sectional brain consist of 30 to 55 coronal slices with 40 µm to 50 µm thickness and 5.1 µm per pixel. Image size varies across brain with resolution ranging from 1 megapixels (1000 *×*1000) to 6 megapixels (2000 *×* 3000) in the sectioning plane. Because the cerebral cortex is easily detached during procedures, if this region is mostly missing in an experimental brain, we preprocess the slices to mask out all the cerebral cortex tissues. The Allen Mouse Brain Atlas is 320 *×* 456 in coronal plane and consists 528 slices, with isotropic 25 µm resolution. For the brain alignment in this systematic anatomical study, we use the brain section in that atlas that contains the region of interest. The cerebral cortex in the atlas is removed with the atlas annotation, if the cerebral cortex in the target experimental brain is mostly missing.

### 4.1. Implementation

#### HOG Cell Size

We used a cell size of 15 pixels to measure image similarity (see Section 3.1.1). This relatively large cell size means we can still capture structural similarity even with uncorrected small distortions. For nonrigid registration in Section 3.2, we decrease the cell size to 4 pixels, because the purpose of this step is to correct distortions. In both steps, the block size is 2 *×* 2. The HOG is computed with the Vlfeat toolbox (Vedaldi and Fulkerson, 2008).

#### Set A

We select a subset A of all slices I to find the best cutting angle and the best corresponding slices. For full brain data which contains about 200 slices, we use about 30 slices with minimal artifacts - 1/6 of the whole sequence. In the anatomical study, each sectional brain consists of about 35 slices. We use every third image for most of the brains - about 12 slices for each experimental brain. For brains with relatively more damaged slices, we manually checked the automatic selection and replaced slices with significant damage with an adjacent good quality slice. Using Matlab, it takes 38.8s on a 12-core 3GHz Linux machine to evaluate a set of 12 slices.

#### Iteration Details

We use three iterations to complete the 2D nonrigid registration described in Section 3.2. The grid spacing is 16 *×* 16 in all iterations. In the first iteration, we downsample both images 4 times in x and y. In the second iteration, images are downsampled by 2*×*. In the final iteration, we operate at the original resolution. The maximum displacement at each level is set to 0.5 times the grid spacing. The actual displacement at the current iteration is the sum of estimated displacements in the previous levels plus the displacement at the current level. The value of data and pairwise terms are computed at the resolution of the current level but projected to the label in the resolution of the final level. Therefore the total number of labels are 113 *×*113. We keep the label set small by only evaluating allowable 17 *×*17 labels (others assigned as a very large number) in the current iteration for each node. Both the unary and the pairwise terms are normalized to range [0, 1]. A coefficient of 0.5 and 1.5 is applied on the point distance term in Eq. 3.3 and the pairwise term respectively. These same parameters were used for all volumes in our dataset. This optimization is computed with graph-cut, alpha expansion optimization for multi-label non-submodular energy (Rother et al., 2007; Boykov et al., 2001; Bagon, 2006).

#### Segmentation

We manually masked the aqueduct of our training brains and downsampled both the experimental slices and aqueduct masks to 512 *×*512. We first labeled five brains (brains were randomly selected in the dataset; one of them is in the evaluated brains) and trained our model on the data-augmented training data. We predicted aqueduct of other brains with the trained model and manually corrected them if necessary. The segmentation network has 9 layers. The input and output image has dimensionality 512 *×*512 *×*1 where the input is the image to be segmented, and the output image is the predicted mask of the aqueduct. The first seven layers have dimension 512 *×*512 *×*32 with dilation rate doubling the rate of the previous layer starting from 1. The convolution kernel size is 3 *×*3. The largest receptive field in the network is the seventh layer - 256 *×*256. The last two layers consist of an undilated smooth layer with the same kernel size and a linear transformation layer. We use intersection over union as the loss function and take 8 points on each half of the segmented aqueduct to compute the point distance term in the revised similarity term function in Eq.19.

### 4.2. Methodology

We use 5 brains in the anatomical study to quantitatively analyze the performance of our framework. In addition, because of the lack of ground truth in these brains, we generated a simulated brain from the atlas with known transformation. The transformation applied partially mimics the property of the real experimental data including different sectioning angle, nonrigid deformation in the sectioning plane, and contrast adjustment.

The most common metric for evaluating image registration is the target registration error (TRE) measured as the Euclidean distance between landmark point coordinates in the target image mapped by the computed registration to the source image and the landmark points in the source image. We asked a neuroscientist to identify 20 sparsely-scattered landmarks in the hind brain of the atlas which she would be confident in locating in both simulated and real experimental brains. The points are enough to cover all the significant brain areas in this study, because 1) an experimental brain has around 35 slices, 2) on a representative experimental slice, there are roughly 30 nuclei identified by its anatomical properties based on neuroscientitsts’ historical consesus, 3) almost all the nuclei are shown at least on 5 slices. The corresponding points of these 20 points are marked by the same neuroscientist in the brains that we evaluated.

In the simulated brain, we can compute the true error of both our method and of the expert, since we have ground truth, as well as the TRE - expert and computation combined error. This information can help interpret the results in the five experimental brains, where we can only compute the TRE.

### 4.3. Comparison Experiment

To compare our results to previous work, we chose to use Ju’s method (Ju et al.,

2006) to reconstruct the brain sections because it is a fully automatic method that contains nonrigid deformation and generates a smooth reconstructed volume which facilitates 3D registration to the reference volume. In previous work, experimental volumes are at least partially reconstructed, because our experimental brains are sectional brains - about 1/6 length of a full mouse brain, we reconstruct the whole section. We first rigidly align the slices and then nonrigidly reconstruct the sections with a five-slice neighborhood. The reconstructed volume is then registered to the reference atlas with Elastix (Klein et al., 2010; Shamonin et al., 2014) and the parameter file in (Hammelrath et al., 2016). This parameter file is developed specifically for 3D mouse brain registration and performs the best among the files that we tested. Reconstructed volumes are first rigidly aligned, affinely aligned, and finally elastically aligned with B-spline to the reference atlas. Similarly with the primary experiment, we measure computation error on the simulated brain and the TRE on both simulated brain and experimental brains. We will show that 3D nonrigid registration is not enough to correct errors introduced by reconstruction.

### 4.4. Results

#### Experiments on Simulated Brain

Figure 7 reports the results of separately measured expert error, computation error, and the TRE - combined expert and computation error - on the simulated brain. The expert has an intrinsic error of about 9 pixels - similar to the TRE of our method. This figure also shows that our method, with a 2.4 pixel error, is about three times better than the reconstruction first approach which has a error of 7.45 pixels.

**Figure 7:**
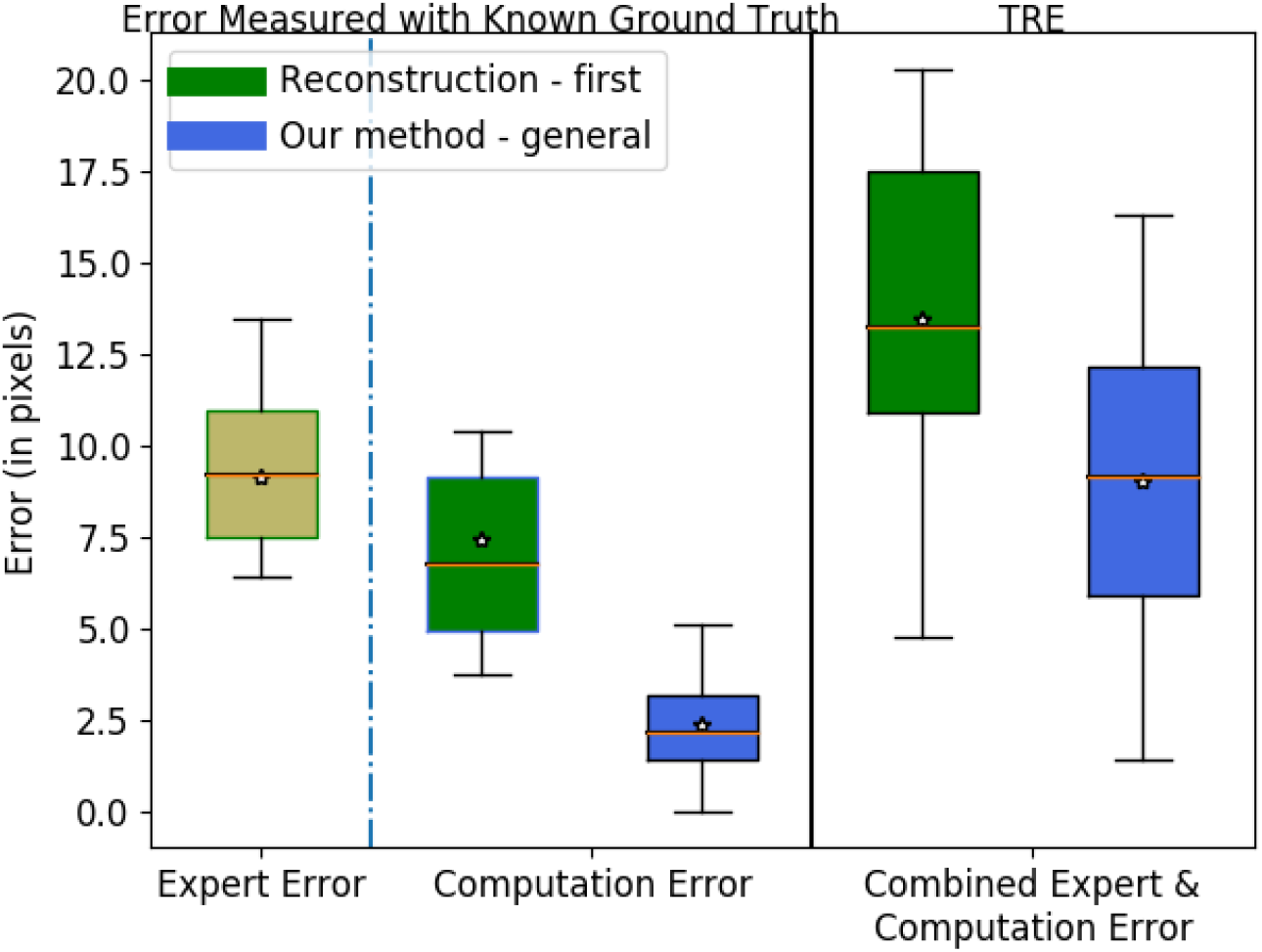
Boxplots showing intrinsic expert error, computation error of the comparison experiment and our method measured with known ground truth, and the TRE -expert and computation combined error - of the reconstruction-first method and our method. The lines on the boxes represent the minimum, first quartile, median (red), third quartile, and maximum respectively. The star denotes the average.

#### Experiments on Real Dataset

Figure 8 reports the experiment results with a reconstruction-first method and our method without and with the data-specific improvement. The five brains represent some data variability we see in the real dataset. These distances represent a combination of human and computer inaccuracy. Based on the simulated result, we believe the intrinsic expert error is likely to be much larger than the computation error. The TRE ratio is on average 2.3 between the reconstruction-first method and our method.

**Figure 8:**
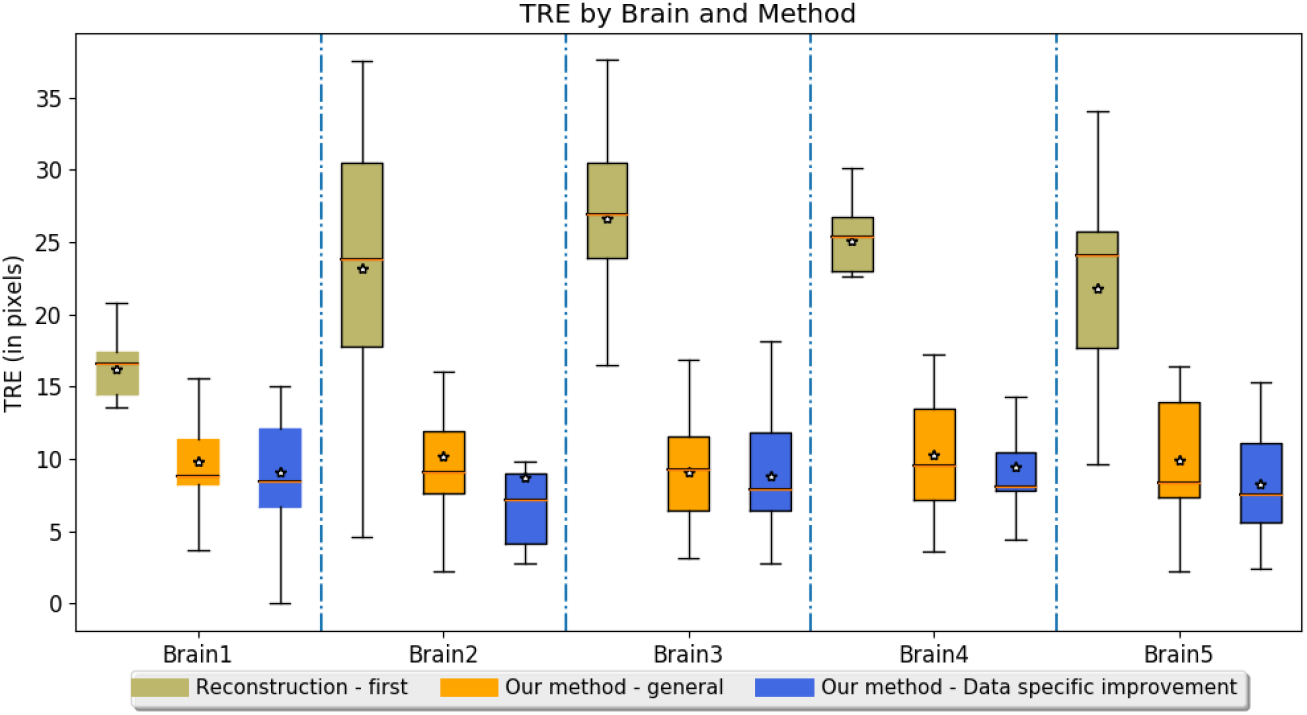
Boxplots of the TRE on evaluated experimental brains.

## 5. Conclusion

Histological sectioning is the most commonly used method to investigate organizations of normal and diseased brains. Individual brain variations, distortions and intensity inconsistency caused by sample preparations make aligning histological brain slices to an atlas a challenging task for both experts and vision algorithms. To address these challenges, we put together a direct approach to solve the mapping problem between a 2D histological sequence and a reference volume which allows us to determine the best corresponding slice for each experimental slice before attempting any nonrigid alignment. It uses the average matching index difference between half-images to create a robust sectioning angle measurement and the HOG L2 norm as the image compar-ison metric. The HOG metric enables image similarity comparison without the need of nonrigid registration. This produces a robust framework that leverages brain structural characteristics and symmetry to determine the cutting angle and matching slices without initial reconstruction which avoids introducing errors as demonstrated by our comparison experiments.

In 2D nonrigid registration, we augmented the standard MRF on medical image registration to model accumulated tension when deforming tissues to more naturally deal with the easily-deformed cavities throughout the brain. This requires us to use squared distance pairwise term, which also required us to pass the simulated stress across iterations. Together our method provides better accuracy to the difficult problem of mapping histological slices to a reference, demonstrating a significant advance over current procedures.

Our approach eliminates the z-shift problem, and works well on both full brain data and sectional data, even for datasets where many slices are corrupted. Since its accuracy is similar to an expert neuroscientist, we have successfully used our method to map multiple brain datasets to the Allen Mouse Brain Atlas, making multi-brain data analysis possible and accurate.

## 6. Acknowledgement

This work is supported by the Hughes Collaborative Innovation Award and a BRAIN initiative grant. We thank Qifeng Chen for suggestions on segmentation networks and donating a GPU for our research, Steven Bell for discussion, proofreading and hardware maintenance, Allen Institute for Brain Science for the reference atlas and Terri Gilbert for the advice on using the atlas.

